# Excitatory and inhibitory D-serine binding to the NMDA receptor

**DOI:** 10.1101/2022.03.07.483247

**Authors:** Remy A. Yovanno, Tsung Han Chou, Sarah J. Brantley, Hiro Furukawa, Albert Y. Lau

**Author notes:** To whom correspondence should be addressed (A.Y.L.), (H.F.).

## Abstract

N-methyl-D-aspartate receptors (NMDARs) uniquely require binding of two different neurotransmitter agonists for synaptic transmission. D-serine and glycine bind to one subunit, GluN1, while glutamate binds to the other, GluN2. These agonists bind to the receptor’s bi-lobed ligand-binding domains (LBDs), which close around the agonist during receptor activation. To better understand the unexplored mechanisms by which D-serine contributes to receptor activation, we performed multi-microsecond molecular dynamics simulations of the GluN1/GluN2A LBD dimer with free D-serine and glutamate agonists. Surprisingly, we observed D-serine binding to both GluN1 and GluN2A LBDs, suggesting that D-serine competes with glutamate for binding to GluN2A. This mechanism is confirmed by our electrophysiology experiments, which show that D-serine is indeed inhibitory at high concentrations. Although free energy calculations indicate that D-serine stabilizes the closed GluN2A LBD, its inhibitory behavior suggests that it either does not remain bound long enough or does not generate sufficient force for ion channel gating. We developed a workflow using pathway similarity analysis to identify groups of residues working together to promote binding. These conformation-dependent pathways were not significantly impacted by the presence of N-linked glycans, which act primarily by interacting with the LBD bottom lobe to stabilize the closed LBD.

## INTRODUCTION

The N-methyl-D-aspartate receptor (NMDAR) is an ionotropic glutamate receptor (iGluR) that uniquely requires the binding of a co-agonist in addition to its primary agonist for activation. This heterotetrameric ion channel comprises at least two different subunits, GluN1 (isoforms 1-4a and 1-4b) and GluN2 (subtypes A-D), assembled as a dimer of GluN1/GluN2 heterodimers. The GluN2 subunit binds the neurotransmitter glutamate, while the GluN1 subunit can either bind the co-agonists glycine or D-serine. Traditionally, glycine had been considered the dominant GluN1 agonist [1–3], but more recent work has suggested that D-serine may in fact be the dominant co-agonist for synaptic NMDARs in the brain [4]. D-serine is synthesized by the enzyme serine racemase expressed in astroglia [5] and neurons [6] [7] and is released into the postsynapse by the Asc-1 transporter [8] [9]. D-serine binding to these synaptic NMDARs is responsible for inducing long-term potentiation (LTP), which is critical for memory functions [10]. In addition, recent clinical efforts have indicated that D-serine could be a promising therapeutic for the treatment of neuropsychiatric disorders [11][12], most notably schizophrenia [13] and post-traumatic stress disorder (PTSD) [14]. Unlike the more well-studied agonists glutamate and glycine, the role of D-serine is less defined, causing it to be known as the “shape-shifting” agonist [9] that can adopt different roles in neurotransmission.

Each NMDAR subunit consists of an amino-terminal domain (ATD), a ligand-binding domain (LBD; also called an agonist-binding domain, ABD), a transmembrane domain (TMD), and a disordered cytoplasmic C-terminal domain [15]. The LBDs adopt a bi-lobed clamshell architecture that close upon agonist binding. Previous computational studies of NMDAR LBDs have indicated that glycine binding to the GluN1 LBD and glutamate binding to the GluN2A LBD drives the conformational equilibrium toward the closed LBD [16]. While crystallographic studies have determined the binding pose of D-serine bound to the closed GluN1 LBD [17], the molecular mechanisms by which D-serine finds its way into and stabilizes NMDAR LBDs is not well understood.

Previous simulation studies have revealed the mechanisms by which glycine and glutamate diffuse into the LBD binding site [18]. Specifically, they found that glycine binds to the GluN1 subunit by freely diffusing into the binding pocket, where it is trapped by energetically favorable interactions with key binding site residues. Glutamate, on the other hand, was found to contact residues along the protein surface that helped guide itself into its binding pocket, positioning it to interact stably with residues in the binding site. These two binding mechanisms were referred to as “unguided" and "guided" diffusion, respectively [19]. This paradigm established the two extremes by which ligands enter their receptor sites: one in which stable ligand binding only depends upon the identity of the binding site residues and another that also heavily relies on residues outside the binding site to guide the ligand toward its bound pose.

Performing multi-microsecond molecular dynamics simulations of the glycosylated GluN1/GluN2A LBD dimer, we identified binding mechanisms and residues critical for promoting D-serine binding and stabilization by developing a new binding pathway clustering workflow. Surprisingly, we observed D-serine binding to both GluN1 and GluN2A LBDs. We determined that D-serine binding to GluN2A partially stabilizes the active LBD conformation. Inspired by these simulation results, we determined that D-serine competes with glutamate for binding to GluN2A via a competitive inhibition mechanism using electrophysiology measurements, where D-serine was found to be inhibitory at high concentrations. Since NMDAR LBDs are glycosylated under physiological conditions [20], including N-linked glycans in our simulations revealed that glycans primarily regulate the binding process by stabilizing the active LBD. In total, we investigated the molecular components contributing to D-serine binding and stabilization, highlighting the complex components driving neurotransmission.

## RESULTS

### D-serine binding pathways for GluN2A and GluN1 LBDs

In simulating the GluN1/GluN2A LBD dimer, which is a physiological NMDAR unit, we intended to focus our attention on the mechanisms by which D-serine binds to the GluN1 LBD, the subunit to which D-serine is a potent agonist. However, in our simulations, we also observed a significant number of D-serine binding events involving the GluN2A LBD, an unexpected finding. These binding events are primarily made up of guided-diffusion pathways in which D-serine contacts key residues on the LBD surface to help guide it into the binding cleft. In our aggregate ~51 *μ*s of sampling of the glycosylated GluN1/GluN2A LBD dimer, we identified 99 guided-diffusion pathways for GluN2A and 104 (plus 23 free diffusion events) for GluN1. Due to the stochastic nature of these pathways, we needed to develop a reliable way to identify key features of predominant binding pathways. To address this, we applied pathway similarity analysis (PSA) [21] to quantify the spatial and geometric similarity between pairs of paths **(Fig. 1A)**. Here, we extend this application to ligand binding pathways by monitoring the change in ligand *C_α_* position throughout each path. This allowed us to cluster paths traversing similar regions of the LBD surface. To aid in describing the different faces of the LBD, we use an order parameter (*ξ*_1_, *ξ*_2_) defined in previous work [16] to describe whether D-serine primarily contacts residues on the *ξ*_1_ or *ξ*_2_ face of the LBD **(Fig. 1B, 2A)**. For GluN2A, cluster analysis revealed four distinct regions of D-serine occupancy. The clusters correspond to the following methods of binding: 1. D-serine approaches the binding pocket from the *ξ*_2_ face; 2. D-serine contacts the D1 residues on the *ξ*_1_ face; 3. D-serine zigzags between D1 and D2 lobes on the *ξ*_1_ face; 4. D-serine primarily contacts residues on the D2 lobe of the *ξ*_1_ face **(Fig. 1C-F)**. Similarly, for GluN1, cluster analysis revealed four distinct clusters corresponding to similar pathways of binding: 1. D-serine contacts the *ξ*_2_ face; 2. D-serine zigzags between D1 and D2 lobes on the *ξ*_1_ face; 3. D-serine contacts residues on the N-terminal (top) end of D1 of the *ξ*_1_ face; 4. D-serine contacts residues of D1 loop 2 that protrudes from the LBD into solution. We then analyzed the resulting clusters to identify key residues that guide D-serine into the binding site **(Fig. 2B-E)**. Interestingly, we observed that GluN1 pathways involve fewer interactions between D-serine and D2 residues; most notably, there were fewer contacts with Helix F (Helix E for GluN2A) compared to GluN2A pathways.

**Fig. 1.**
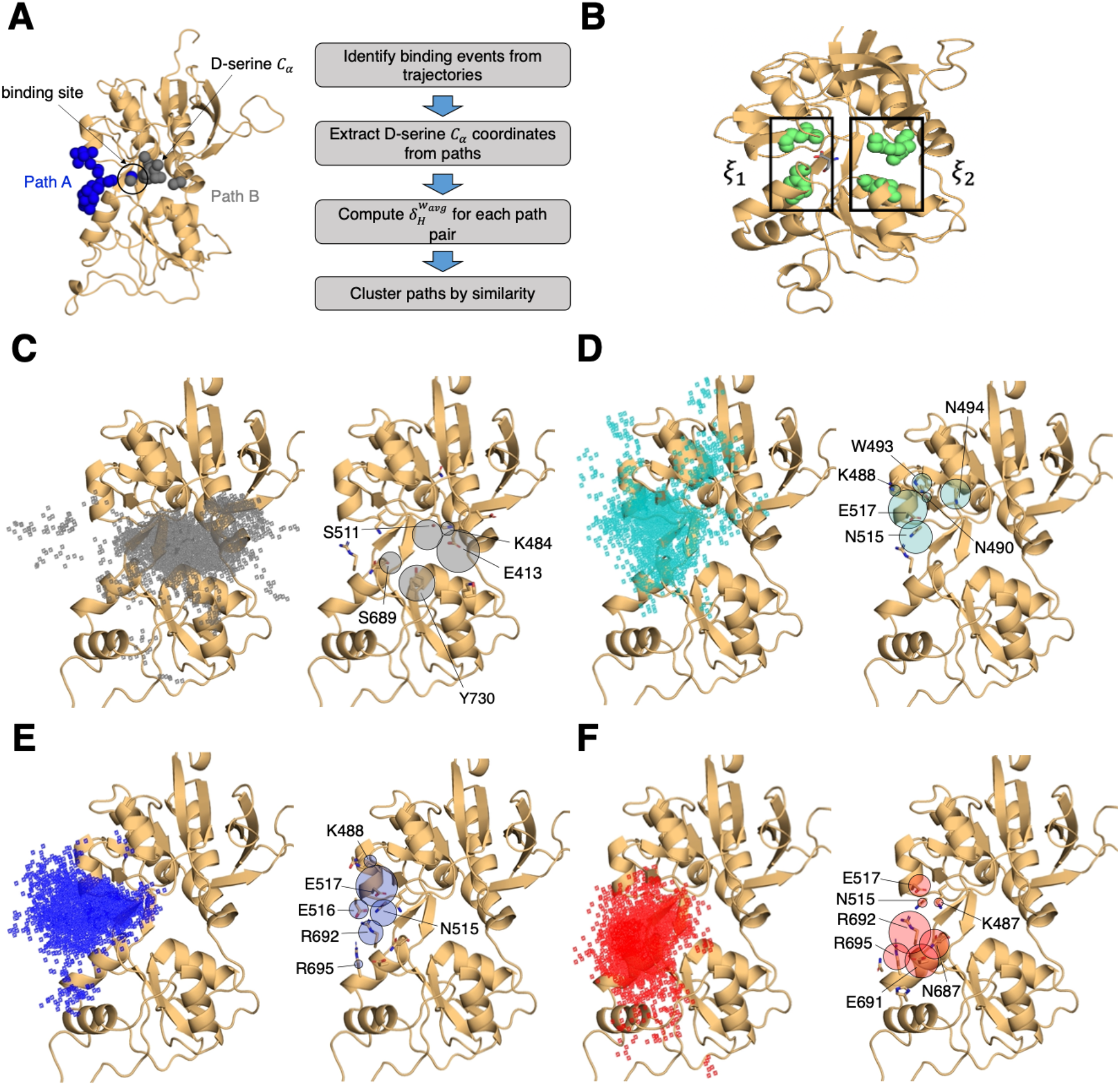
Identifying D-serine binding pathways for GluN2A using pathway similarity analysis (PSA). **(A)** Overview of the PSA workflow for quantifying similarity between D-serine binding pathways. **(B)** 2-dimensional order parameter (*ξ*_1_, *ξ*_2_) that describes the degree of GluN2A LBD closure. For each of the above **(C-F)**, the left image shows D-serine density, while the right image shows the residues most frequently contacted by D-serine as it enters/leaves the binding site for each cluster. Labeled residues demonstrate ≥ 0.2 fractional occurrence defined relative to the most contacted residue in each cluster, but all residues with ≥ 0.1 fractional occurrence are shown in stick representation (see **Dataset S5**). **(C)** Cluster 1 involves residues of the *ξ*_2_ face of the LBD. **(D)** Cluster 2 involves residues of the *ξ*_1_ face of the D1 lobe. **(E)** In Cluster 3, D-serine zigzags between D1 and D2 lobe residues of the *ξ*_1_ face. **(F)** Cluster 4 primarily involves D2 lobe residues on the *ξ*_1_ face.

**Fig. 2.**
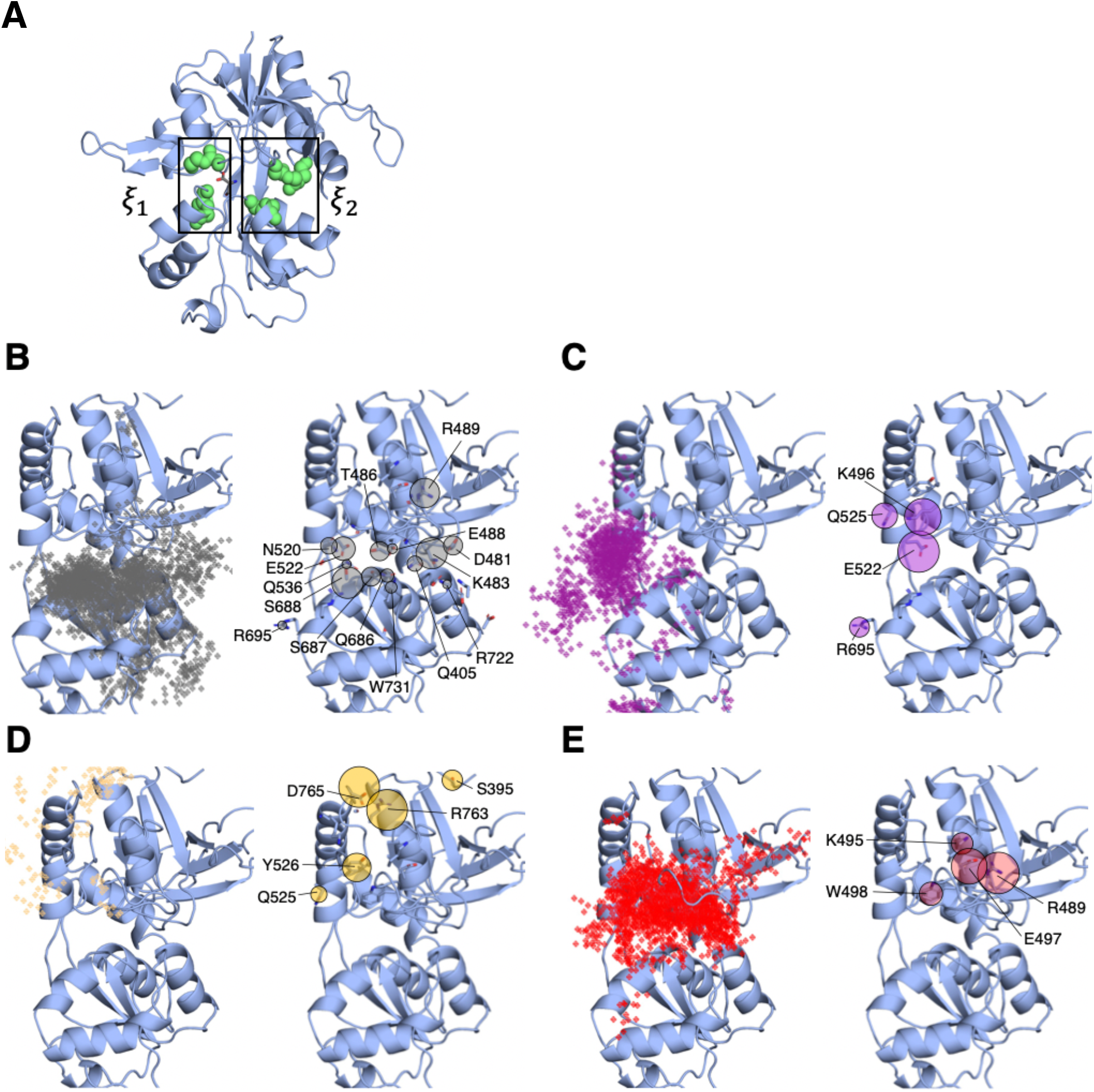
Identifying D-serine binding pathways for GluN1 using pathway similarity analysis (PSA). **(A)** 2-dimensional order parameter (*ξ*_1_, *ξ*_2_) that describes the degree of GluN1 LBD closure. For each of the above **(B-E)**, the left image shows D-serine density, while the right image shows the residues most frequently contacted by D-serine as it enters/leaves the binding site for each cluster. Labeled residues demonstrate ≥ 0.2 fractional occurrence defined relative to the most contacted residue in each cluster, but all residues with ≥ 0.1 fractional occurrence are shown in stick representation (see **Dataset S6**). **(B)** In Cluster 1, D-Serine contacts residues on the *ξ*_2_ face of the LBD. **(C)** Cluster 2 involves interactions with both D1 and D2 residues of the *ξ*_1_ face. **(D)** Cluster 3 involves contacts with residues at the top of the D1 lobe on the *ξ*_1_ face. **(E)** Cluster 4 is defined by interactions with D1 loop 2 that reaches into solution.

To quantify the extent to which these clusters involve similar residue contacts, we used a pairwise similarity metric called the overlap coefficient (i.e., Szymkiewicz– Simpson coefficient) that describes agreement between sets of residues [22]. Doing so provides a way to determine whether these spatial clusters are mostly made up of random contacts, or whether groups of residues tend to act together to promote binding, allowing us to quantify the extent to which agonist diffusion is “guided” by contacts along the LBD. For GluN2A, we computed the overlap coefficient for all path pairs in each cluster for comparison with the global mean (global ⟨*OC*⟩ = 0.557) **(Fig. S1A)**. We found that pathway pairs in three of the four clusters yielded an overlap coefficient greater than the mean of all pairs of paths from all clusters, indicating that pathways in each cluster are made up of specific residue contacts **(Fig. S1C)**. In contrast, for GluN1, a significant cluster (26 paths) involving interactions with residues on the *ξ*_2_ face of the LBD has a cluster mean *OC* much less than the global mean (global ⟨*OC*⟩ = 0.671), indicating that this cluster primarily comprises random contacts **(Fig. 1B, S1B,D)**. This suggests that D-serine binding to GluN1 may be more diffusion-driven and less guided than to GluN2A. Therefore, we propose that agonist binding mechanisms exist on a spectrum ranging from unguided to guided diffusion. The difference in the specificity of D-serine contacts along binding pathways for GluN2A and GluN1 suggests that the extent to which agonists rely on pathways of guiding residues depends on LBD architecture and not solely upon the identity of the agonist.

Mapping important pathway residues onto the intact GluN1/GluN2A NMDAR (PDB ID: 6MMM [23]) further enriches our understanding of binding pathways by allowing us to determine whether residues in particular pathways are accessible for binding or obscured by other receptor domains and subunits. For GluN2A, access to residues on the extreme of the *ξ*_2_ face is slightly restricted by the presence of the GluN1 subunit of the adjacent LBD dimer **(Fig. S2A)**. However, this interface does not seem to be near the specific residues identified as critical for binding. Even more restricted is access to residues on the *ξ*_1_ face of GluN1, which are obscured by GluN2A of the adjacent LBD dimer, including residues identified as critical for binding pathways **(Fig. S2B)**. This might bias the pathways observed for the intact receptor by forcing the agonist to favor residues on the *ξ*_2_ face of the LBD. Since our overlap coefficient analysis of the cluster that corresponds to the *ξ*_2_ face of GluN1 identified more non-specific interactions, it is possible that the D-serine mechanism would be biased to favor unguided diffusion. It is also possible that access to residues near the N-terminal end of D1 would be restricted by the R2 lobe of its own ATD.

We next investigated whether a specific LBD conformational state was favored for successful D-serine binding pathways. We computed our (*ξ*_1_, *ξ*_2_) order parameter to quantify the degree of closure of the LBDs for all trajectory frames identified as part of binding (and unbinding) pathways and found that (*ξ*_1_, *ξ*_2_) = (16,14) for GluN2A **(Fig. S3A)** and (*ξ*_1_, *ξ*_2_) = (11,13) for GluN1 **(Fig. S3B)**. These values correspond to a partially open LBD. The LBD needs to be open enough for the ligand to diffuse into the pocket but closed enough to form some stabilizing interactions with the ligand. However, we notice that the *ξ*_1_ is smaller for GluN1, indicating that agonist binding can occur at slightly more closed LBD conformations. GluN1 pathways where (*ξ*_1_, *ξ*_2_) = (11,13) are mostly in the cluster defined by D-serine interactions with Loop 2, highlighting the role of Loop 2 residues in D-serine binding to GluN1. Overall, these results suggest that the degree of LBD closure does influence the likelihood of successful binding.

### Effects of D-serine binding on the LBD conformational free energy landscapes

Since we did not expect to see D-serine binding to the GluN2A LBD, we needed to determine whether these GluN2A D-serine binding events are able to modulate the GluN2A LBD conformation. Since full LBD closure occurs on multi-microsecond to millisecond timescales [24][25][26], direct observation of such a conformational change was not fully captured from our equilibrium binding trajectories. Instead, to ensure we are sampling the full range of LBD conformations, we performed umbrella sampling free energy molecular dynamics simulations to obtain the conformational free energy landscape of GluN2A bound to D-serine **(Fig. 3A)**. We used the order parameter (*ξ*_1_, *ξ*_2_) [16] that captures the opening and closing motion of the LBDs observed in crystal structures of these domains. Since no crystal structure exists for D-serine bound to GluN2A, we identified residues critical for stabilizing the agonist in the closed state by analyzing contacts in lowest-energy (≤1 kcal mol^-1^) conformers **(Fig. S4A)**. For reference, we compared the resulting energy landscape to those previously computed for the apo- and glutamate-bound GluN2A monomers **(Fig. 3C,D)** [16]. We see that, like glutamate, D-serine stabilizes the closed LBD conformation. The D-serine energy landscape has a global minimum corresponding to (*ξ*_1_, *ξ*_2_) values of (11, 11.5 Å) and a metastable minimum corresponding to (*ξ*_1_, *ξ*_2_) values of (15.5, 11.5 Å). The presence of a metastable agonist-bound LBD partially open intermediate suggests that D-serine may not stabilize the closed conformation to the same extent as glutamate and generate sufficient force to control channel gating. We then compared different conformers corresponding to these two states to determine residues critical for agonist stabilization. The primary difference between the residue contacts in conformers of the two states is the prevalence of interactions with Thr-690 **(Fig. S4B)**, which only contacts D-serine in the more closed state centered at (*ξ*_1_, *ξ*_2_) = (11, 11.5 Å). This is supported by our binding simulations; although we do not fully sample LBD closure, trajectory frames with low (*ξ*_1_, *ξ*_2_) values involve contacts with Thr-690. This suggests that Thr-690 is critically involved in promoting full GluN2A LBD closure upon agonist binding.

**Fig. 3.**
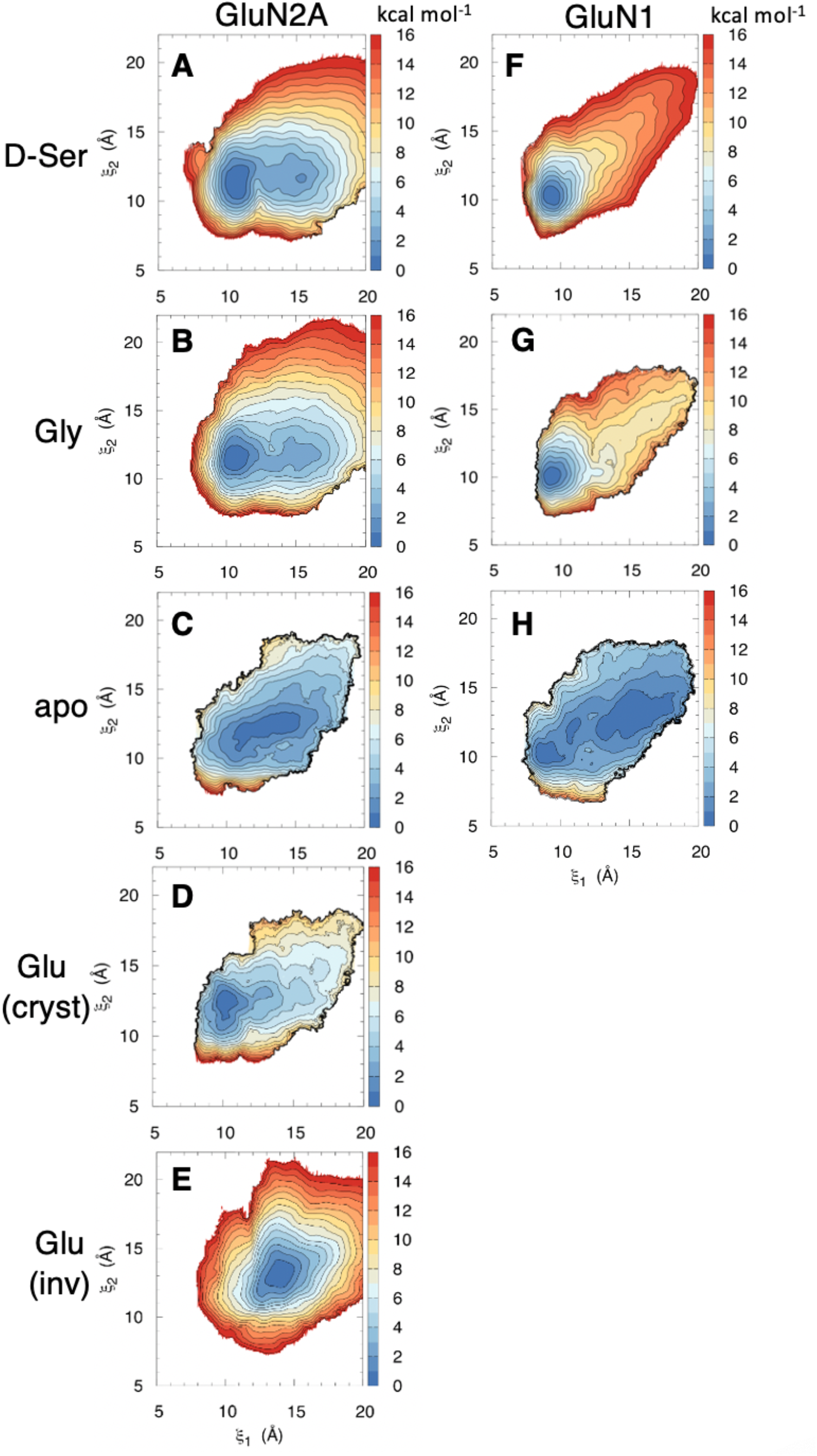
Conformational free energy landscapes for GluN2A and GluN1 LBDs. Umbrella sampling molecular dynamics simulations were used to compute the potential of mean force (PMF) along the (*ξ*_1_, *ξ*_2_) order parameter for **(A)** D-serine bound to GluN2A, **(B)** glycine bound to GluN2A, **(C)** apo GluN2A previously computed in [16], **(D)** glutamate bound to GluN2A in its crystallographic pose previously computed in [16], **(E)** glutamate bound to GluN2A in the inverted pose identified in [18], **(F)** D-serine bound to GluN1, **(G)** glycine bound to GluN1 previously computed in [16], **(H)** apo GluN1 previously computed in [16].

Experimental binding studies have indicated that D-serine may be a more potent GluN1 agonist than glycine [27]. To better understand the molecular mechanism responsible for this difference in agonist potency, we computed the conformational free energy for the D-serine-bound GluN1 LBD **(Fig. 3F)**. Compared with the previously computed glycine-bound and apo LBDs **(Fig. 3G,H)** [16], the presence of D-serine in the binding cleft results in a greater population of conformers in the closed conformation and fewer conformers adopting a more open conformation. Similar to GluN2A Thr-690, GluN1 Asp-732 and (to a lesser extent) Ser-688 help stabilize D-serine in the closed LBD conformation by interacting with the D-serine hydroxyl. For this reason, we propose that D-serine’s high potency is due, at least in part, to its ability to more strongly stabilize a closed LBD through additional interactions with the D2 lobe.

### D-serine and glutamate compete for binding to the GluN2A LBD

Since our simulations revealed that D-serine can enter the GluN2A LBD binding pocket and partially stabilize the active conformation, we hypothesized that D-serine might compete with glutamate for binding to GluN2A. In fact, we observed D-serine binding to GluN2A, even in the presence of glutamate, although glutamate bound more frequently than D-serine and with longer residence times in the binding site **(Datasets S2, S3)**. Since increasing the D-serine concentration would increase the frequency of D-serine binding to GluN2A, it is possible that D-serine could function as an inhibitor (competitive antagonist) at high concentrations. If true, this behavior may factor into therapeutic strategies focused on increasing D-serine concentration in the synapse by establishing an upper dosage limit after which a D-serine increase is no longer potentiating.

To probe this behavior experimentally, we measured GluN1-2A NMDAR currents using two-electrode voltage clamp (TEVC) electrophysiology. We observed that at high (~1 mM) D-serine concentrations, NMDAR activity was inhibited **(Fig. 4A)**. The inhibition was dependent on glutamate concentrations, implying that the inhibitory effect of D-serine may be competitive **(Fig. 4B)**. Furthermore, dose-response curves of glutamate activation were right-shifted in the presence of increasing concentrations of D-serine **(Fig. 4C)**. The calculated slope value of the Schild plot at 1.1 ± 0.1 implied that D-serine and glutamate likely compete against each other **(Fig. 4C)**. Combined with our simulation results, our electrophysiological data supports the hypothesis that D-serine at high concentrations can bind to the GluN2A subunit and compete against glutamate.

**Fig. 4.**
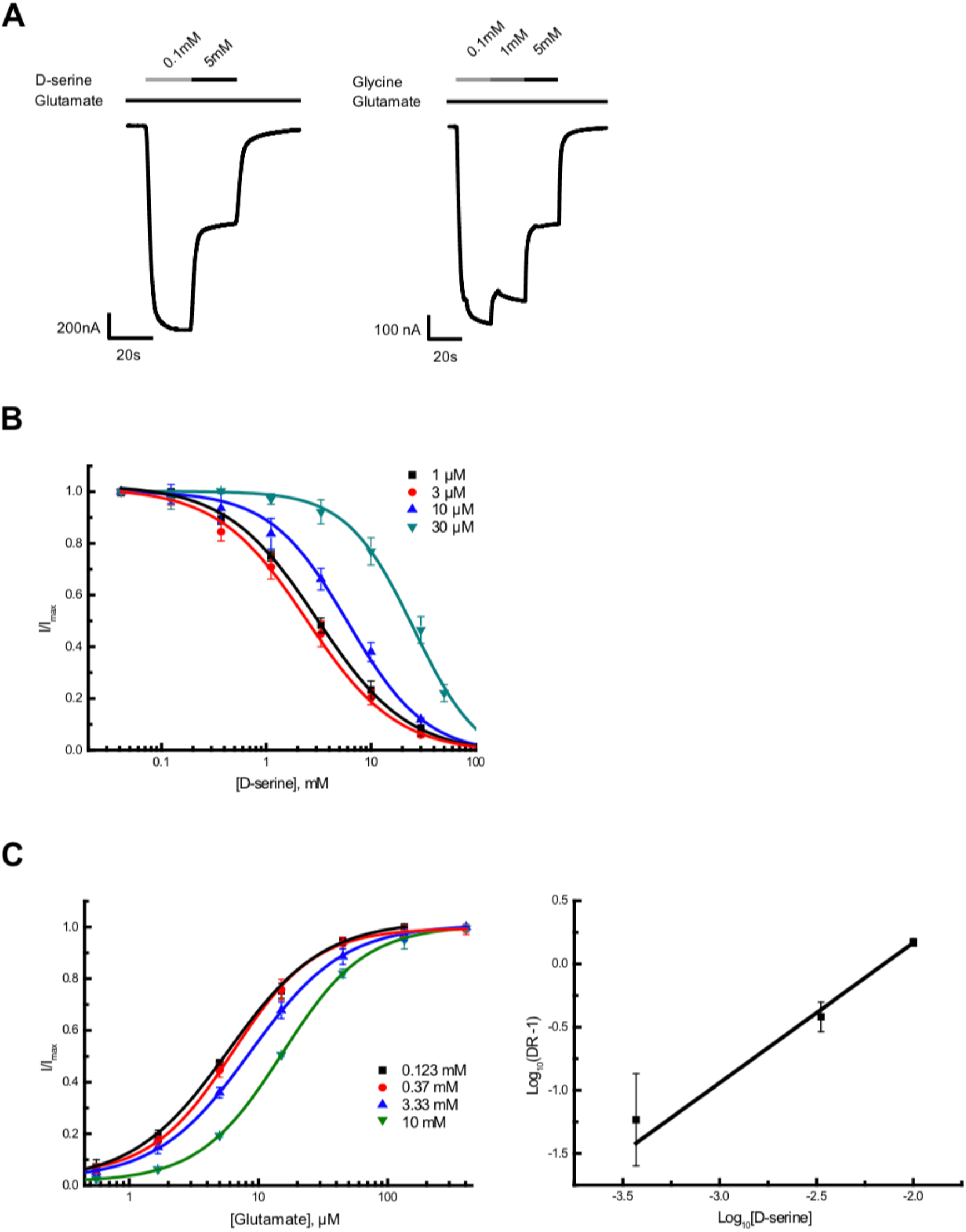
D-serine completes glutamate binding as an antagonist at high concentration. **(A)** Representative Two-electrode voltage clamp (TEVC) recording on GluN1/GluN2A NMDARs expressing oocytes. The trace showed GluN1 agonist D-serine inhibited the NMDAR current at a high concentration. 6 *μ*M of glutamate was present throughout the recording. (**B)** Glutamate-concentration dependent dose-response curves of high-concentration D-serine inhibition. (**C)** D-serine-concentration dependent dose-response curves of glutamate potentiation (left). Schild plot analysis of D-serine competition on glutamate (right). The calculated slope of the Schild plot was 1.11±0.13 and the intercept was 2.38±0.26. DR stands for dose ratio. All the dose-response experiments were repeated at least four times.

Since a similar inhibitory effect was also observed at high glycine concentrations by TEVC electrophysiology **(Fig. 4A)**, we repeated our umbrella sampling simulations with glycine bound to the GluN2A LBD. We see that glycine also favors the closed LBD **(Fig. 3B)**. The lowest-energy conformers of GluN2A with glycine are fastened shut by contacts between the N-terminal amine of glycine and Tyr-730. Although glutamate still stabilizes the closed GluN2A LBD to the greatest extent, comparable thermodynamics between different agonists suggests that kinetics of agonist binding and unbinding is a critical driver of agonist-induced activation. The GluN2A LBD likely never closes around glycine because glycine does not remain bound long enough to induce LBD closure.

Previous binding studies [18] have indicated that glutamate, the primary GluN2A agonist, similarly relies on LBD surface residues to promote binding. To determine whether D-serine and glutamate binding are guided by similar residue contacts, we computed the overlap coefficient between residues in D-serine and glutamate pathways to be 0.964 for the glycosylated GluN2A LBD, corresponding to a significant overlap in agonist occupancy **(Fig. 5A)**. This high degree of overlap between glutamate and D-serine pathway residues indicates that they bind through similar mechanisms.

**Fig. 5.**
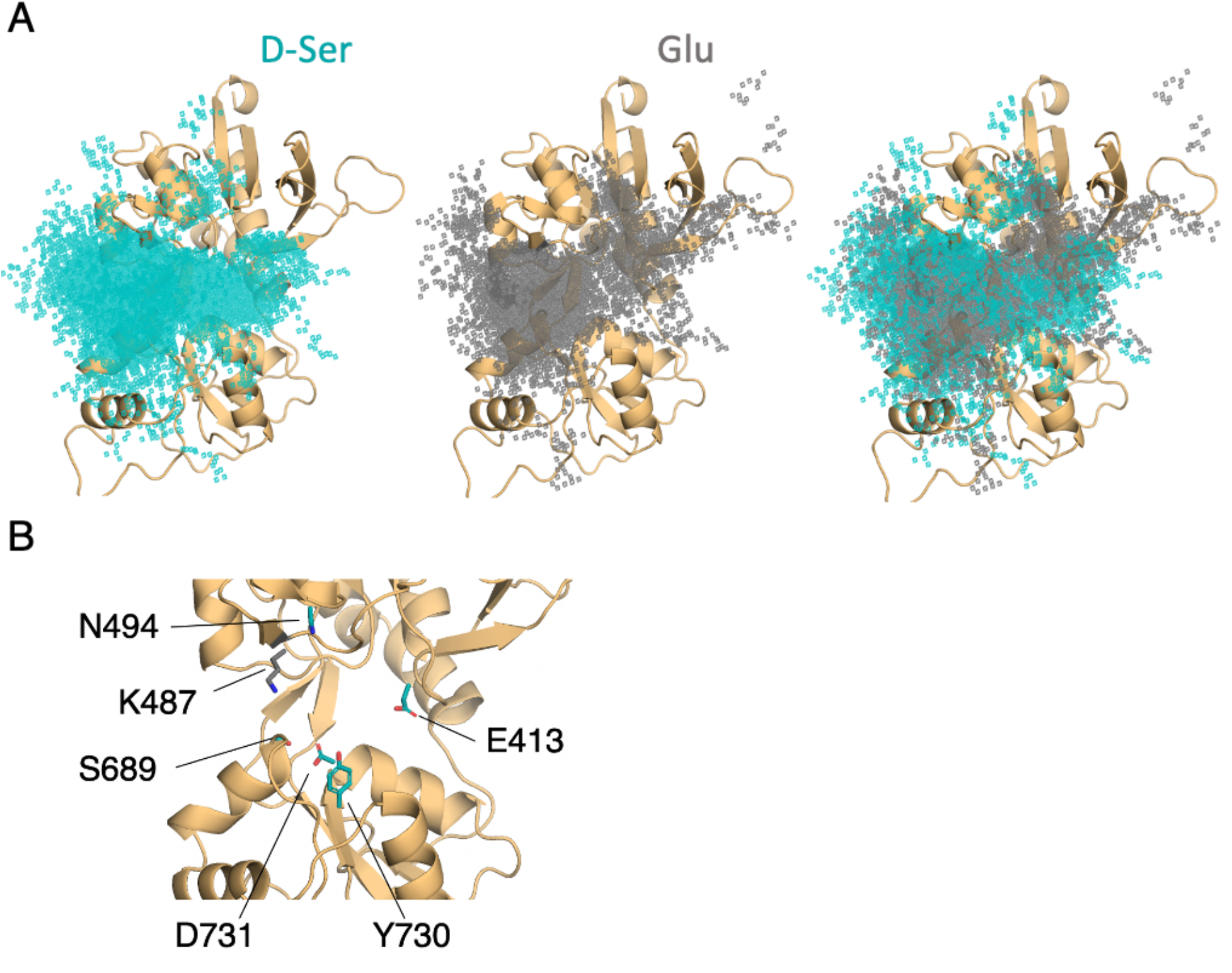
Comparison of D-serine and glutamate binding to GluN2A. **(A)** Overlay of D-serine (teal) and glutamate (gray) density. **(B)** Residues that distinguish D-serine (teal) from glutamate (gray) binding pathways (see **Dataset S7**).

Despite similar pathway residues, we identified key residues that distinguish glutamate from D-serine binding pathways **(Fig. 5B and Dataset S7)**. Most of the residues important for D-serine binding, but not for glutamate binding, are located on the *ξ*_2_ face of the LBD. Most notably, Glu-413, Tyr-730, Ser-511, and Asp-731 all occur in D-serine binding pathways with a frequency of more than ten times their fractional occurrence in glutamate binding pathways. It is important to note, however, that glutamate does interact with residues on the *ξ*_2_ face, but the specific nature of those contacts differ between the two agonists. In contrast, we found that Lys-487 is contacted with significantly greater frequency in glutamate binding pathways. Due to these residues’ close proximity to the binding cleft, it is likely that these residues are responsible for facilitating proper positioning of the agonists in the binding site, based on differences in agonist size and shape.

An important feature of glutamate binding to GluN2A is its ability to bind in an inverted pose relative to the crystal structure, which we observed in previous simulations [18] [19]. Since no experimental structure exists for glutamate bound in the inverted pose, we performed umbrella sampling simulations to determine the free energy landscape of the GluN2A LBD with glutamate bound in the inverted pose **(Fig. 3E)**. We found that glutamate bound in the inverted pose prevents full LBD closure as predicted in previous work [18]. Specifically, glutamate in the inverted pose stabilizes a conformation centered around (*ξ*_1_, *ξ*_2_) values of (14, 13 Å). Comparing the low-energy conformers of D-serine and inverted glutamate (≤ 1 kcal mol^-1^) with the glutamate-bound crystal structure, we found that D-serine and glutamate are stabilized by the same residues, although there are fewer interactions between Thr-690 and glutamate in the inverted pose, further supporting the importance of this residue for stabilizing the fully closed LBD.

### Kinetic analysis of D-serine binding pathways

We computed the D-serine association rate constant (*k_on_*) for GluN2A and GluN1 LBDs using a method described [28] and used in previous iGluR work [19] as summarized in the equation below:

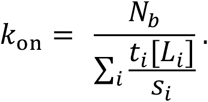

Here, *N_b_* is the number of association events, *t_i_* is the time the agonist spends in bulk solvent, *s_i_* is the number of identical binding sites, and [*L_i_*] is the concentration of free agonist. One advantage of this approach is the ability to combine simulations performed at various concentrations of free agonist [*L_i_*]. Here, *k_on_* is a bulk property and relies on fully sampling the LBD conformational landscape throughout the simulation. However, our binding simulations fail to adequately sample the agonist-bound, closed LBD state. This affects both the number of observed binding events *N_b_* and the time the agonist spends in bulk solvent (*t_i_*). Since this value is most sensitive to the number of identified binding events *N_b_*, we computed the *k_on_* for different *N_b_* values based on the duration of the resulting binding event. This minimizes contributions from extremely short binding events that are unlikely to be functionally relevant. For GluN2A, this results in a D-serine *k_on_* with an upper bound of 7.8 × 10^7^ M^-1^s^-1^ (all events included) and a lower bound of 1.6 × 10^7^ M^-1^s^-1^ (only events with agonist residence times > 100 ns were included). For GluN1, the upper bound for *k_on_* is 9.0 × 10^7^ M^-1^s^-1^ and the lower bound is 7.0 × 10^6^ M^-1^s^-1^. Based on these values, it is reasonable to expect that D-serine binds to GluN2A and GluN1 at similar rates. For comparison, the association rate constants computed for glutamate binding to GluN2A with this method range from 4.9 × 10^7^ M^-1^s^-1^ to 1.4 × 10^8^ M^-1^s^-1^. Similar ranges of D-serine binding rate constants for GluN2A and GluN1 support our data indicating a guided-diffusion mechanism. However, this definition of the association rate constant does not capture the molecular details that produce this bulk behavior.

For agonist binding mechanisms dominated by guided diffusion, we can monitor how much time the agonist spends (1) in bulk solvent, (2) associated with the LBDs, and (3) docked in the binding cleft (interacting with the conserved arginines Arg-523 for GluN1 or Arg-518 for GluN2A). Transitions between these states can be represented by the following three-step process:

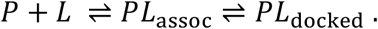

Here, the *PL_assoc_* state either results in successful binding (represented by pathways) or nonspecific interactions resulting in dissociation. From the clusters of residues that we identified in our pathway similarity analysis, we determined to what extent a particular residue is critical for guiding the agonist into the binding site using a conditional probability-based framework **(Datasets S9, S10)**. For GluN2A, given that a binding event results in successful agonist docking, residues Asp-515, Glu-517, Arg-692, Asn-687, Lys-487, Lys-484, and Ser-689, Lys-488, Ser-511, and Glu-413 are contacted most frequently across all datasets. Given successful D-serine binding, contacts with GluN1 residues Lys-496, Lys-495, Trp-498, Arg-489, and Glu-497 occur in the greatest number of pathways. Slightly less agreement in crucial GluN1 binding residues across datasets further supports a more diffusive/random binding mechanism for D-serine binding to GluN1.

### Role of N-linked glycans in D-serine binding pathways

In addition to identifying residues that are responsible for agonist specificity in binding pathways, we also explored the effect of the N-linked Man5GlcNAc2 (Man5) glycans **(Fig. S6A)** on the residues involved in agonist binding pathways. Previous electrophysiological studies have indicated that glycans function as LBD potentiators [29]. In our simulations, we observed that near-pocket glycans appear to “reach” into the binding pocket. This reaching behavior was observed in previous simulations of the glycosylated NMDAR LBDs in which the glycan forms a “cage” around the binding pocket by forming interactions with the LBD D2 lobe and is believed to be associated with NMDAR potentiation by glycans [29]. For GluN2A, there are two glycans that are near the binding pocket: N443-Man5 and N444-Man5, both of which can interact with the LBD D2 lobe **(Fig. 6A)**. For GluN1, there is a single glycan N491-Man5 that adopts this caged conformation **(Fig. 6B)**. To quantify this behavior in our simulations, we developed a general order parameter to describe the relationship between the glycan and the LBD D2 lobe that measured the minimum distance between any glycan heavy atom and any residue on the LBD D2 lobe. From this order parameter, we computed glycan PMFs along the glycan-D2 order parameter for each near-pocket glycan **(Fig. 6C-E)**.

**Fig. 6.**
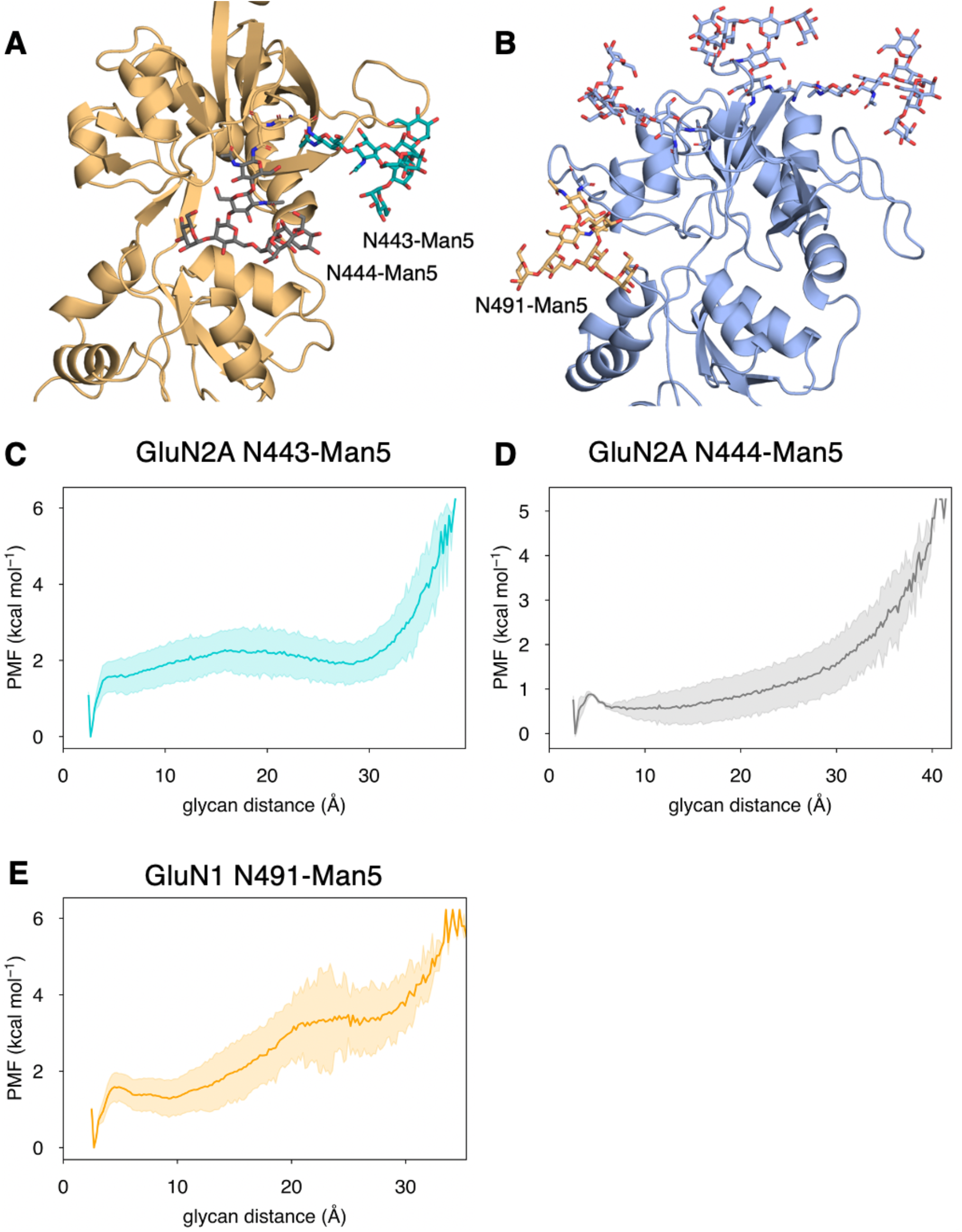
Conformational dynamics of near-pocket glycans. N-linked Man5GlcNAc2 (Man5) glycans **(A)** N443-Man5 and N444-Man5 for GluN2A and **(B)** N491-Man5 for GluN1. Glycan conformational energy landscapes for **(C)** GluN2A N443-Man5, **(D)** GluN2A N444-Man5, and **(E)** GluN1 N491-Man5 were obtained by computing the minimum distance between all glycan heavy atoms and D2 lobe residues and binning the distribution from all glycosylated simulation systems. Shaded error regions were computed using a block-averaging scheme described in Methods.

We compared our glycosylated trajectories with an additional 30 *μs* of simulations of the non-glycosylated GluN1/GluN2A LBD dimer to identify ways in which the presence of glycans influence binding pathways. Our data indicate that residues on the *ξ*_2_ face are contacted more frequently in non-glycosylated simulations, although these residues are important for D-serine binding with and without glycans **(Dataset S11)**. GluN2A residues Asp-515 and Glu-517, are contacted more frequently in glycosylated systems. The frequency with which D-serine interacts with GluN1 residue Arg-489 in pathways is greater for glycosylated pathways than those without glycans. On average, glycan-mediated D-serine interactions result in slightly longer pathways, suggesting that the presence of glycans slows down the binding process, setting up small kinetic “traps”.

When we analyzed glycan behavior in our binding pathways, we found that very few D-serine binding pathways (27% for both GluN2A and GluN1) involve contacts with glycans. While glycan-agonist interactions make up a small percentage of time spent in binding pathways (10% for GluN2A and GluN1 D-serine pathways), patterns in glycan interactions with the agonist as it binds suggest that glycans contribute to binding pathways in a consistent way. The most common glycan-mediated D-serine-LBD interactions for GluN2A involve an interaction network formed by N443-Man5 with Glu-412, Lys-438 **(Fig. S6B)**, Lys-738, Glu-413 **(Fig. S6C)**, Tyr-730, and Ser-511 **(Fig. S6D)**, as D-serine moves into the binding pocket. Another contact network formed by N444-Man5 with Lys-487, Asn-687 **(Fig. S6E)**, Arg-692, Arg-695, **(Fig. S6F)**, and Glu-413 (alongside the N443-Man5 glycan). For GluN1, the N491-Man5 glycan interacts with D-serine, trapping it in a network of interactions dominated by Arg-489 **(Fig. S6G)**. When formed, this contact network functions as a kinetic trap that results in longer binding pathways. Additionally, the N440-Man5 glycan also contacts D-serine as it interacts with Arg-489 and Glu-497 **(Fig S6H)**. It is interesting to note that, unlike the glycan-mediated contacts identified for GluN2A, glycan-mediated agonist contacts for GluN1 do not involve D2 lobe residues. These glycan-mediated interactions illustrate how glycan conformation can play a functional role through involvement with agonist binding and LBD conformational dynamics. However, since glycan-mediated interactions are so infrequent, the potentiating effect of glycan-D2 interactions dominates functionally.

We quantified the dependence of glycan conformation on agonist binding and LBD conformation by comparing glycan PMFs for different LBD conformations. For GluN2A, we found that glycan-D2 interactions occur more readily when the LBD is closed (calculated using a 1-dimensional projection of our LBD order parameter *ξ*_12_, see Methods). This effect was more dramatic for N443-Man5 than for N444-Man5 **(Fig. S7A,B)**. A similar relationship was determined for the N491-Man5 glycan of GluN1 **(Fig. S7C)**; this is consistent with previous simulations [29] that suggest that N491-Man5 acts as a latch that stabilizes LBD closure. No significant relationship between glycan-D2 distance and the presence of an agonist (D-serine, glutamate, or both) in the binding site was observed.

## DISCUSSION

Here, we characterized the guided-diffusion mechanism that drives D-serine binding to NMDAR LBDs. Instead of binding solely to the GluN1 LBD, we observed substantial D-serine binding to the GluN2A LBD, a subunit widely accepted to bind to the neurotransmitter glutamate. We showed by electrophysiology that D-serine at high concentration can compete against glutamate at GluN2A, which in turn inhibits the channel activity. In the context of synaptic transmission, our finding implies that D-serine could play a role in modulating the strength of synaptic transmission. The synaptic concentration of glutamate ranges from nanomolar concentrations [30] to >1 mM following an action potential [31]. The synaptic concentration of D-serine is unclear, however; the extracellular concentration of D-serine ranges from 5 to 7 μM [32] [33]. Possible routes for D-serine to enter the synapse include vesicular release by astroglia [34] and transport by Asc-1 [35].

Free energy landscapes computed for GluN2A bound to glutamate [16], D-serine, and glycine all indicate stabilization of the closed LBD bi-lobe, which is the conformational state required for receptor activation. Agonists that can interact extensively with bottom-lobe residues stabilize this state. Since glutamate does this to the greatest extent, it is likely that D-serine does not generate sufficient force to fully gate the ion channel. Subtle differences in the thermodynamics of agonist stabilization suggest that kinetics further distinguish individual agonists. While glutamate has a slightly higher association rate than D-serine, differences between association rates across agonists and subunits is not drastic. We hypothesize that, in order for agonist binding to result in NMDAR activation, the agonist must remain in the binding site long enough to induce closure – we found that this is largely dependent upon the number and strength of stable contacts the agonist forms with both D1 and D2 lobe residues.

We determined the role of N-linked glycans in agonist binding and stabilization. Glycans impact agonist binding kinetics less by direct glycan-agonist interactions and more by stabilizing the closed LBD through glycan-D2 interactions. This bias toward LBD closure would increase the agonist residence time and potentiate NMDAR activity.

Our adaptation of pathway similarity analysis allowed us to identify clusters of residues critical for binding agonists. This also allowed us to determine that the presence of pathways depends on the degree of LBD closure. We also observed that D-serine binds to GluN2A using similar pathways and residues as glutamate, while the locations of key D-serine cluster residues for GluN1 are different. Applied more broadly to drug-binding simulations, this method of analyzing binding pathways provides a useful framework for gleaning biological insight from noisy and diffusive binding data.

## METHODS

### Equilibrium Molecular Dynamics Simulations

A construct of the GluN1/GluN2A dimer based on crystal structure (PDB ID: 2A5T [36]) used in our previous study [18] was used as a starting model. The residue numberings are based on the Uniprot numbering for GRIN1 and GRIN2a entries. Man5GlcNAc2 (Man5) glycans were added using CHARMM-GUI *Glycan Reader & Modeler* [37] [38] [39] [40] to asparagine residues 440, 471, 491, and 771 of GluN1 and asparagine residues 443 and 444 of GluN2A in accordance with physiologically relevant glycosylation sites [20]. GluN2A was chosen as the GluN2 subtype both to facilitate comparison with previous simulation studies and because recent evidence has suggested that the GluN2A subtype is the primary subtype at synapses, where D-serine is the dominant co-agonist [4].

All systems were solvated in a 140 Å *×* 110 Å *×* 110 Å orthorhombic water box with ~150 mM NaCl using CHARMM [41]. All systems were electrically neutral. All simulations in this work were performed using the CHARMM36 forcefield [42] and TIP3P water model [43]. The systems were pre-equilibrated using NAMD 2.13 [44] first using NVT conditions and gradually relaxing backbone-sidechain restraints and then for 15 ns using NPT conditions at a pressure of 1 atm and a temperature of 310 K. The pre-equilibrated systems were then simulated on Anton 2 provided by the Pittsburgh Supercomputer Center [45]. A weak center-of-mass restraint of 0.5 kcal mol^-1^ Å^-1^ was applied to GluN2A N, CA, and C atoms of residues 461-463. 507-509, and 523-525 to prevent large protein translational motion. Simulations on Anton 2 were carried out at 310 K with the NPT ensemble and with the weak center-of-mass restraint of 0.3 kcal mol^-1^ Å^-1^ in accordance with previous simulations [19]. Additional simulation details are provided in **Dataset S1**.

### Identification of binding pathways

Identifying frames in which the ligand is bound in the receptor’s binding pocket provides key information about the ligand’s binding affinity and the bound ensemble; however, it fails to account for the process by which the ligand enters and leaves the binding pocket. In guided diffusion, the residues that guide the ligand into the binding pocket are critical for promoting the bound state. While imposing a simple distance cutoff is sufficient for identifying the fully bound state, identifying the pathways by which the ligand binds is less trivial. Here, we introduce a “binding chains” paradigm for defining the ligand’s path along the protein. These binding chains are defined from ligand association to dissociation. An association begins when any polar ligand heavy atom comes within 6 Å of any protein polar heavy atom. The ligand is considered associated until is diffuses beyond 10 Å from the protein. The resulting chains are then filtered by contact with the selected “docking” residue(s). Here, we use the conserved arginine residue for each subunit (Arg-523 for GluN1 and Arg-518 for GluN2A) as the essential docking residue. These chains are filtered then split into their “binding” and “unbinding” components by a more specific docking criterion. In our case, we require that the NH1 and NH2 atoms of the conserved arginine be within 4 Å of the ligand carboxyl in accordance with the following scheme:

Condition 1: Arg NH1 is within 4 Å of the ligand OT1 <u>AND</u> Arg NH2 is within 4 Å of the ligand OT2 <u>OR</u>
Condition 2: Arg NH2 is within 4 Å of the ligand OT1 <u>AND</u> Arg NH1 is within 4 Å of the ligand OT2

This scheme accounts for both the crystallographic binding pose (Condition 2) and a “flipped” ligand orientation (Condition 1). Chains that fail to meet these criteria are discarded. Since binding and unbinding pathways can be considered reversible, we combine them in our analysis, reversing the order of the unbinding pathways so that all pathways have the same directionality. This results in a series of binding pathways we can characterize both geometrically and in terms of key residue interactions.

### Pathway Similarity Analysis and Clustering

Pathway Similarity Analysis (PSA) was applied to each binding pathway by monitoring the agonist position as it binds. PSA involves computing a pairwise distance metric between paths that serves as a measure of geometric similarity [21]. The weighted average Hausdorff distance was selected as the path metric because it gave the most geospatially distinct clusters of agonist density around the protein. This weighted average Hausdorff distance was computed for all pairs of paths using the following formula as described in previous work [21] and implemented in the MDAnalysis python package [46][47]. The weighted-average Hausdorff distance between two paths *A* and *B* can be expressed as:

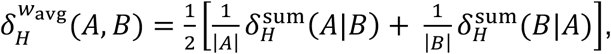

where |*A*| and |*B*| are the number of frames in paths *A* and *B*, respectively, and 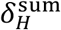 is the one-sided summed Hausdorff distance from path *A* to path *B*,

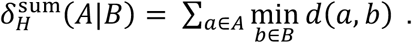

Here, *d*(*a*, *b*) represents the distance between point *a* of path *A* and point *b* in path *B*.

For our system, each point *a* is the agonist *C_α_* position for a single frame in path *A*, and each point *b* is the agonist *C_α_* position for a single frame in path *B*. Therefore, *d*(*a*, *b*) represents the Euclidean distance between the agonist *C_α_*’s of points in paths *A* and *B*. 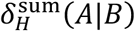 is then computed by summing the shortest distance from each point *a* in path *A* to any point *b* of path *B* over all points in path *A*. Each of the normalized one-sided sums 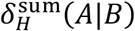 and 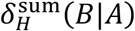 are then averaged with equal weights. This does not give more weight to pathways with more frames, thus removing the temporal component from the analysis. Temporal patterns in binding pathways are analyzed for the spatial clusters separately.

These path pairs were then clustered using hierarchical clustering according to their weighted-average Hausdorff distances with the Ward (minimum variance) linkage criterion as described in previous work [21] and implemented in SciPy [48]. The complete linkage criterion also gave reasonable clustering. This agglomerative metric assigns clusters by successively combining clusters that minimize the sum of squared errors between them. Hierarchical clustering presents an advantage here because it does not assume the number of clusters *a priori*. Rather, final clusters were selected using the Ward distances showed in the dendrograms (see supplemental) as a guide and by overlaying the ligand occupancy density on the protein to ensure that each cluster represents a distinct spatial region of the protein.

### Quantifying residue similarity with the overlap coefficient (Szymkiewicz–Simpson coefficient)

To quantify the similarity between two sets of residues *A* and *B*, the overlap coefficient was computed by dividing the number of overlapping residues between *A* and *B* by the size of the smaller set of residues and is illustrated in the equation below [22]:

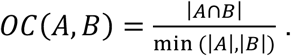

Scaling the size of the intersection by the smallest set size normalizes the overlap and accounts for the large range in pathway lengths. If *A* is a subset of *B*, then *OC*(*A*, *B*) = 1. This scaling method is appropriate, since these pathways are stochastic and involve a mixture of random residue contacts and “guiding” residue contacts critical for binding. This would be problematic for the more common Jaccard similarity metric, which scales the intersection by the total size of both sets, where many random contacts increase pathway length and dilute the value of the similarity metric.

The overlap coefficient was used to quantify the residue overlap between pairs of pathways in each cluster to validate the spatial clustering and determine whether pathways within clusters involve similar residue contacts. In addition, this metric was used to quantify the similarity between residues involved in D-serine and glutamate binding.

### Umbrella Sampling

All-atom models were constructed from monomeric GluN1 (PDB ID: 1PB8 [17]) and GluN2A (based on PDB ID: 2A5S [36]). Since no crystal structure of D-serine bound GluN2A exists, LBDs were constructed using MODELLER [49] to fill in missing residues, and sidechain remodeling was performed on those residues using SCWRL4 [50]. D-serine and glycine were modelled into the GluN2A LBD by superimposing the conserved arginine of the 2A5S glutamate-bound crystal structure (Arg-518) with the conserved arginine of the D-serine (1PB8) or glycine (1PB7) bound crystal structure, since there exists no crystal structure of GluN2A bound to these agonists. Bound crystallographic waters in the GluN2A (2A5S) and GluN1 (1PB8) structures were retained in the simulations.

To generate windows for umbrella sampling, targeted molecular dynamics simulations were performed by “opening” the closed LBD along the order parameter (*ξ*_1_, *ξ*_2_) [16]. Specifically, *"* and are defined as the center of mass distance between the backbone atoms of the following residue selections: **ξ*_1_* is defined by residues 484-485 and 688-689 for GluN1 and residues 485-486 and 689-690 for GluN2A. *ξ*_2_ is defined by residues 405-407 and 714-715 for GluN1 and 413-414 and 713-714 for GluN2A. 205 simulation windows were selected at 1 Å × 1 Å increments. Each window was solvated with a solvent box with dimensions 94 Å × 72 Å × 68 Å and 150 mM NaCl.

Umbrella sampling simulations were performed by applying a bias of 2 kcal mol^-1^ to the (*ξ*_1_, *ξ*_2_) order parameter to each of the 205 simulation windows. Equilibration was performed in an NVT ensemble by gradually relaxing backbone and sidechain restraints, and production simulations were carried out in an NPT ensemble at 300 K and 1 atm for best comparison with previously computed NMDAR LBD monomers [16]. To ensure that the agonist does not diffuse out of the binding site, a restraint of 2 kcal mol^-1^ Å^-1^ between the carboxyl group of the agonist and the guanidinium group of the conserved arginine (Arg-523 for GluN1 and Arg-518 for GluN2A) was applied if the distance between these groups exceeded 3.2 Å. Previous work has indicated that these restraints do not affect the results but ensures that only the bound population is sampled [16]. A weak center-of-mass restraint of 0.5 kcal mol^-1^ Å^-1^ was used applied to the N, CA, and C atoms of residues 461-463, 507-509, and 523-525 for GluN2A and residues 460-462, 512-514, and 528-530 for GluN1 to prevent translational protein motion. Biased trajectories were mathematically unbiased using the weighted histogram analysis method (WHAM) [51] [52]. 5 ns of production sampling for each window were used to compute the potential of mean force (PMF) for each simulation agonist. Standard deviations of all PMFs were computed by block averaging with ten blocks of trajectory for each window [53].

### Computing energetics of glycan conformational dynamics

To quantify glycan conformational dynamics, a glycan-D2 order parameter was defined as the minimum distance between the heavy atoms of the glycans near the binding cleft (N491-Man5 for GluN1 and N443-Man5 and N444-Man5 for GluN2A) and the bottom lobe *C_α_* atoms (residues 537-544 and 663-754 for GluN1 and residues 533-539 and 661-757 for GluN2A). One relative PMF was computed for each of the three near-pocket glycans using a window size of 0.2 Å using all glycosylated datasets. Error for each PMF was quantified using the standard deviation computed by block averaging with five blocks **(Fig. S5A-D)**. Blocks for which the window is not sampled were omitted from the error calculation; this was only necessary for high glycan distances >20 Å. A 1D projection of the (*ξ*_1_, *ξ*_2_) order parameter, *ξ*_12_, which averages *ξ*_1_ and *ξ*_2_, was used as a single measure of LBD closure for computing glycan PMFs [16][54][55].

### Electrophysiology

cRNA encoding GluN1-4b and GluN2A was injected into defolliculated *Xenopus laevis* oocytes (0.2–0.5 ng total cRNA per oocyte). The oocytes were incubated in recovery medium (50% L-15 medium (Hyclone) buffered by 15mM Na-HEPES at a final pH of 7.4), supplemented with 100 μg mL^-1^ streptomycin, and 100 U mL^-1^ penicillin at 18°C. Two electrode voltage clamp (TEVC; Axoclamp-2B) recording was performed between 24 to 48 hours after injection using an extracellular solution containing 5 mM HEPES, 100 mM NaCl, 0.3 mM BaCl_2_, 10mM Tricine at final pH 7.4 (adjusted with KOH). The current was measured using agarose-tipped microelectrode (0.4–0.9 MΩ) at the holding potential of −60 mV. Maximal response currents were evoked by 50 μM of D-serine and 100 μM of L-glutamate. Data was acquired by the program PatchMaster (HEKA) and analyzed by Origin 8 (OriginLab Corp).

## Acknowledgements

Anton 2 computer time (MCB130045P) was provided by the Pittsburgh Supercomputing Center (PSC) through NIH grant R01GM116961 (to A.Y.L.); the Anton 2 machine at PSC was generously made available by D.E. Shaw Research. We also used resources provided by the Maryland Advanced Research Computing Center (MARCC) at Johns Hopkins University. This work was funded by the Johns Hopkins Catalyst Award (to A.Y.L.); NIH T32GM135131 (to R.A.Y. and S.J.B.); NIH NS111745 and MH085926 (to H.F.); Robertson funds at CSHL, Doug Fox Alzheimer’s fund, Austin’s purpose, Heartfelt Wing Alzheimer’s fund, and the Gertrude and Louis Feil Family Trust (to H.F.).

## Author Contributions

R.A.Y. conducted molecular dynamics simulations; R.A.Y., S.J.B., and A.Y.L. analyzed the results; T.-H.C. and H.F. designed and conducted experiments related to electrophysiology; R.A.Y., T.-H.C., H.F., and A.Y.L. wrote the manuscript.

