## Supplemental Information for "Excitatory and inhibitory D-serine binding to the NMDA receptor"

[*] To whom correspondence should be addressed:

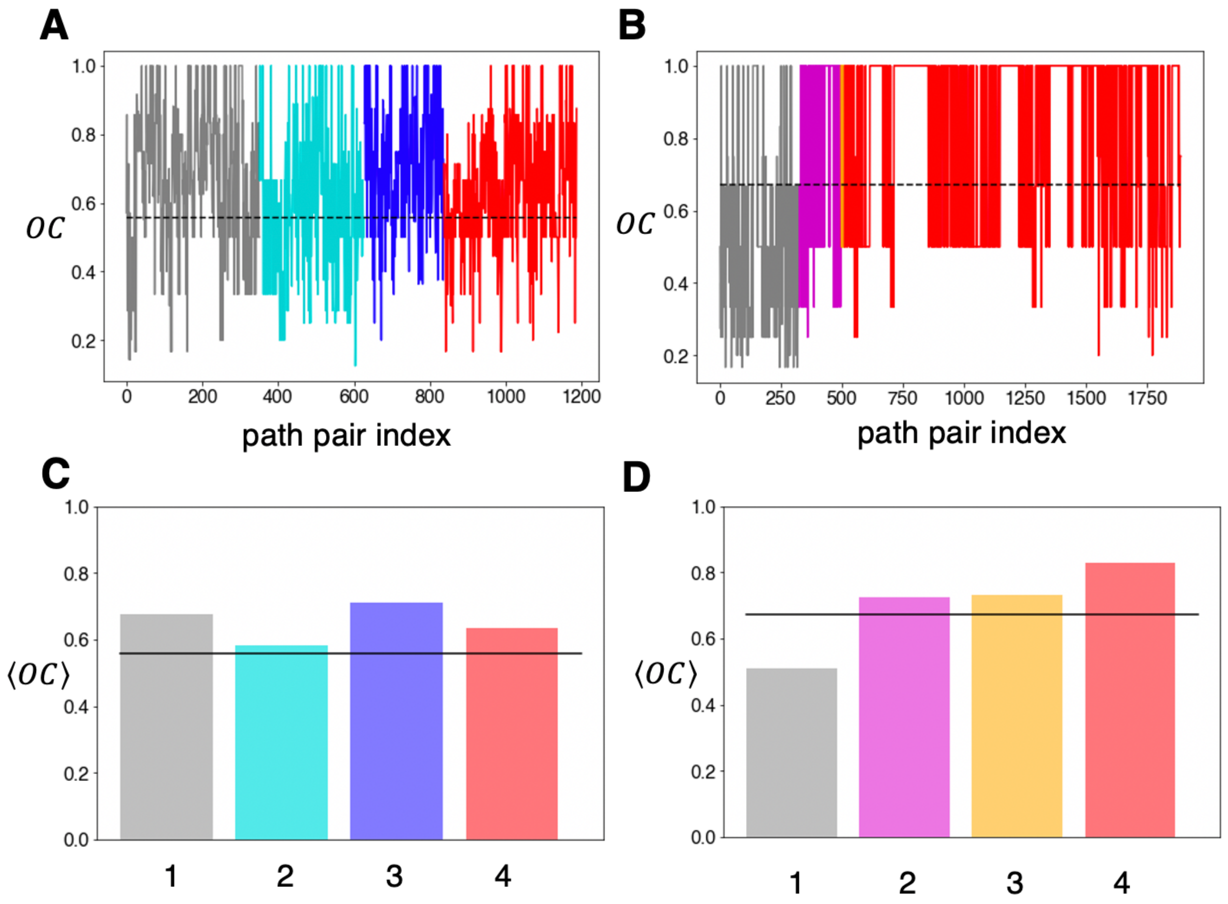


**Figures 1, 2 – figure supplement 1.** Overlap coefficient analysis for GluN2A and GluN1 binding pathways. The overlap coefficient was computed for each pair of paths within each cluster for **(A)** GluN2A and **(B)** GluN1. The dotted line in each plot indicates the global mean $\langle OC \rangle$ = 0.557 for GluN2A and $\langle OC \rangle$ = 0.671 for GluN1. The mean $OC$ for each cluster was computed for **(C)** GluN2A and **(D)** GluN1. For reference, the black solid line indicates the global mean $OC$.

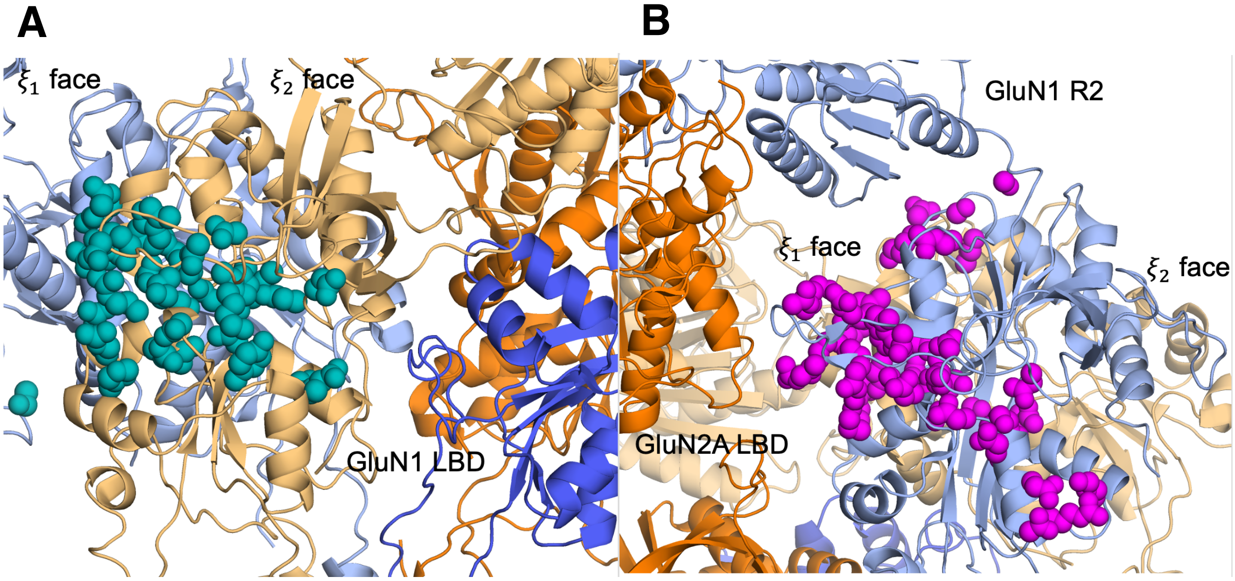


**Figures 1, 2 – figure supplement 2.** D-serine pathway residues mapped onto the intact GluN2A NMDAR (PDB ID: 6MMM[1]). Sidechain atoms are shown as spheres for **(A)** GluN2A in teal and **(B)** GluN1 in magenta and the adjacent domains labelled.

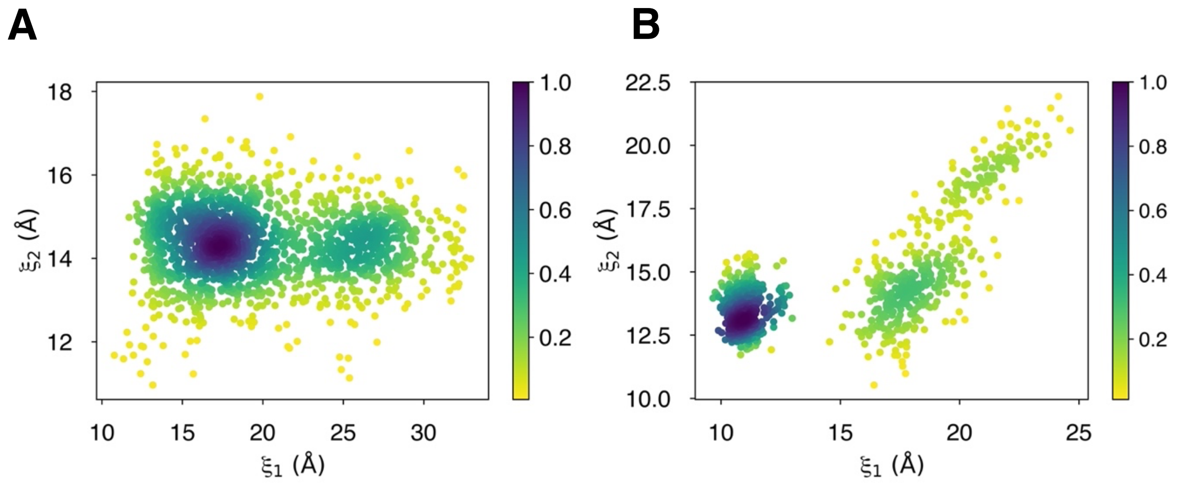


**Figures 1, 2 – figure supplement 3.** Degree of LBD closure during D-serine binding pathways to **(A)** GluN2A and **(B)** GluN1. The two-dimensional $(\xi_1, \xi_2)$ order parameter was computed for all frames in binding/unbinding pathways and was plotted as a function of relative density as indicated by the colorbar.

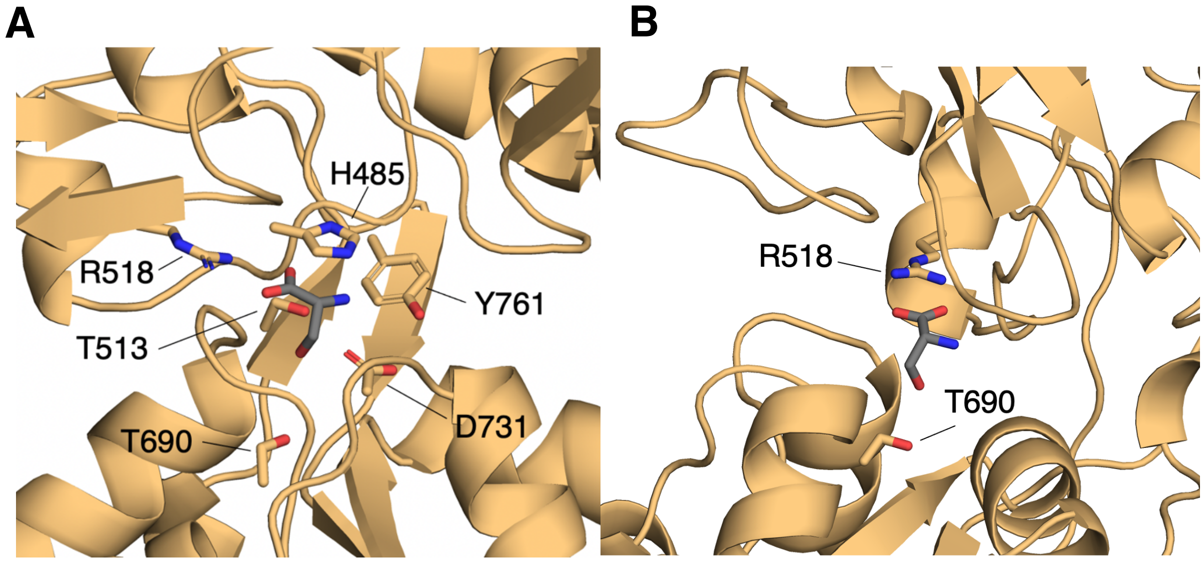


**Figure 3 – figure supplement 1. (A)** Binding-site residues for D-serine bound to the GluN2A LBD computed from lowest-energy conformers (≤ 1 kcal $mol^{-1}$) from umbrella sampling simulations. Residues with >50% contact frequency in the ensemble of lowest-energy conformers are labelled here. **(B)** D-serine interaction with Thr-690 present in lowest-energy conformers.

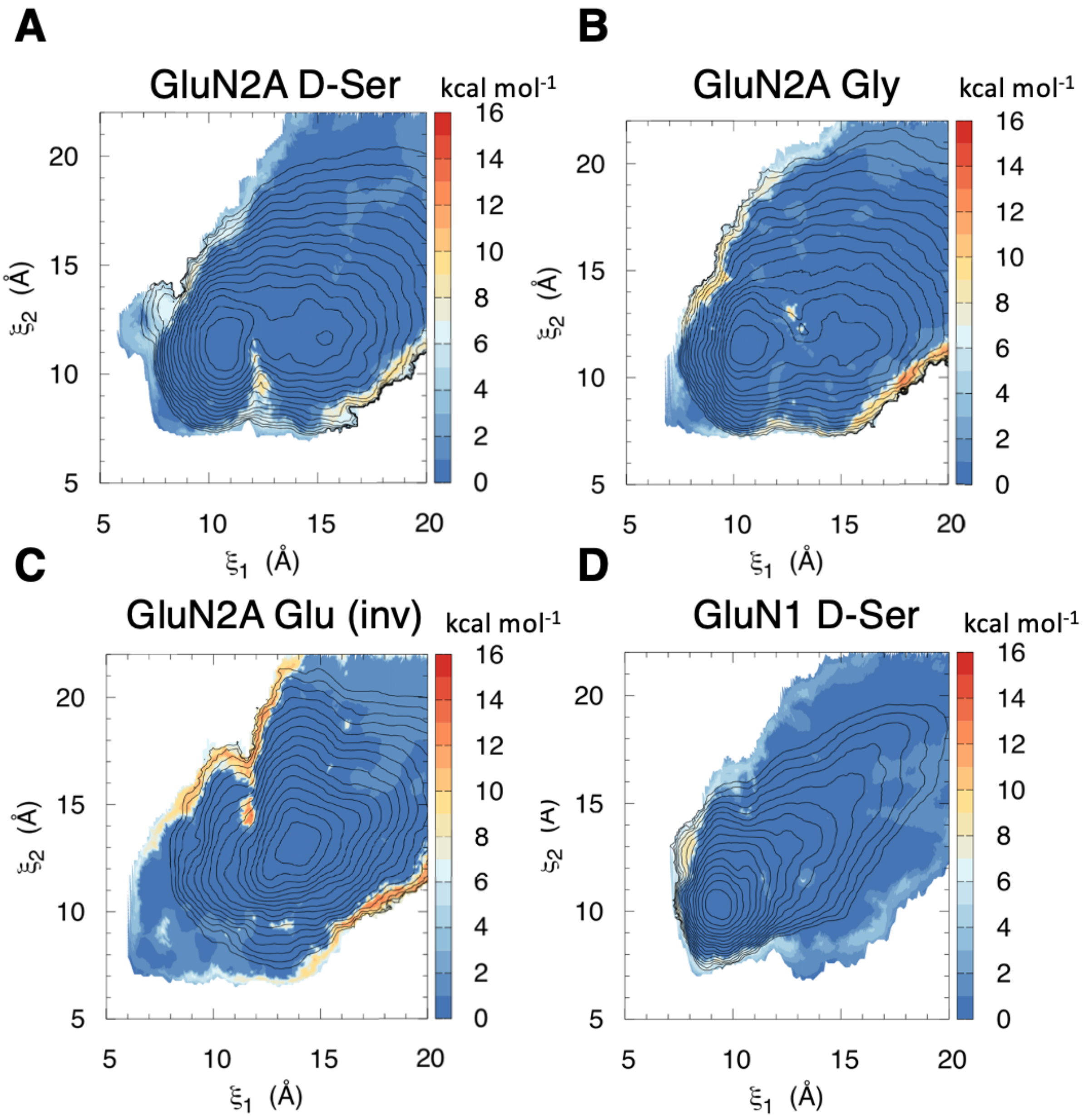


**Figure 3 – figure supplement 2.** Error of umbrella sampling PMFs computed by block averaging for **(A)** D-serine bound to GluN2A, **(B)** glycine bound to GluN2A, **(C)** glutamate bound to GluN2A in the inverted pose, and **(D)** D-serine bound to GluN1. The colors of the colorbar correspond to the standard deviation in kcal $mol^{-1}$, which was computed for each window over 10 blocks.

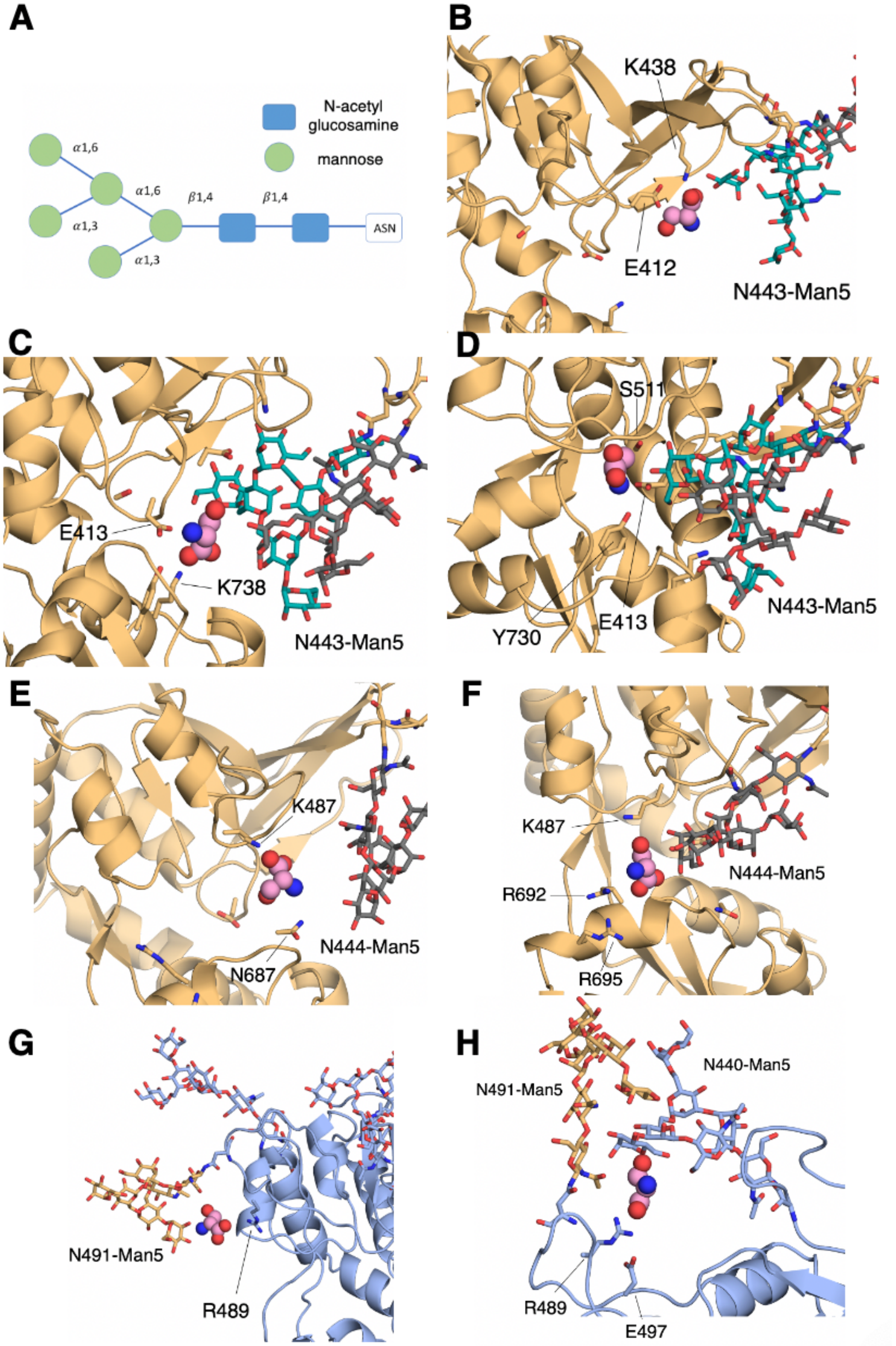


**Figure 6 – figure supplement 1.** N-linked Man$_5$GlcNAc$_2$ (Man5) glycans interacts with D-serine as it binds. **(A)** Schematic of the Man5 glycan. **(B)** Contact network involving the GluN2A N443-Man5 glycan and residues Glu-412 and, Lys-438, **(C)** Lys-738 and Glu-413, **(D)** Glu-413,Tyr-730, and Ser-511. **(E)** Contact network involving the GluN2A N444-Man5 glycan and residues Lys-487 and Asn-687, **(F)** Lys-487, Arg-692, and Arg-695. **(G)** Contact network formed between D-serine, the GluN1 N491-Man5 glycan, and Arg-489. **(H)** Additional GluN1 contact network formed between both GluN1 N491-Man5 and N440-Man5 glycans, residues Arg-489 and Glu-497, and D-serine.

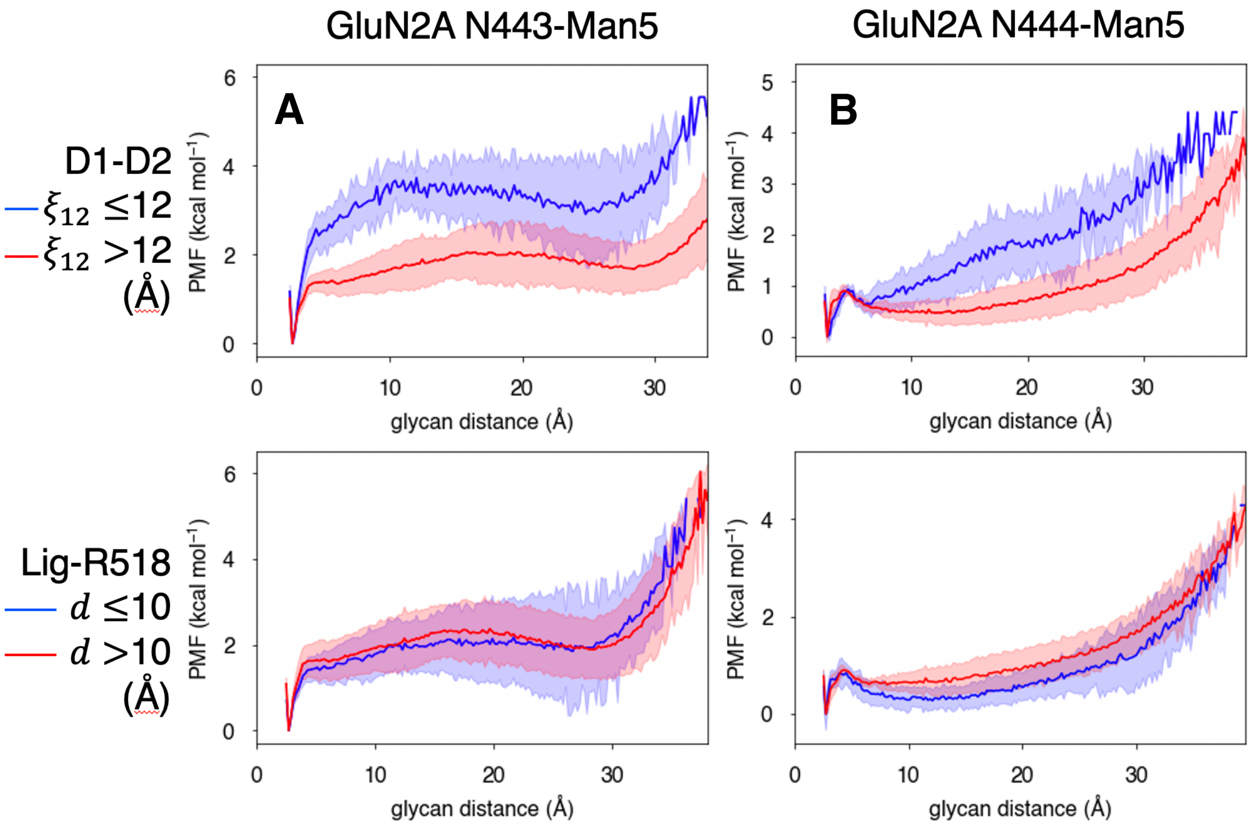


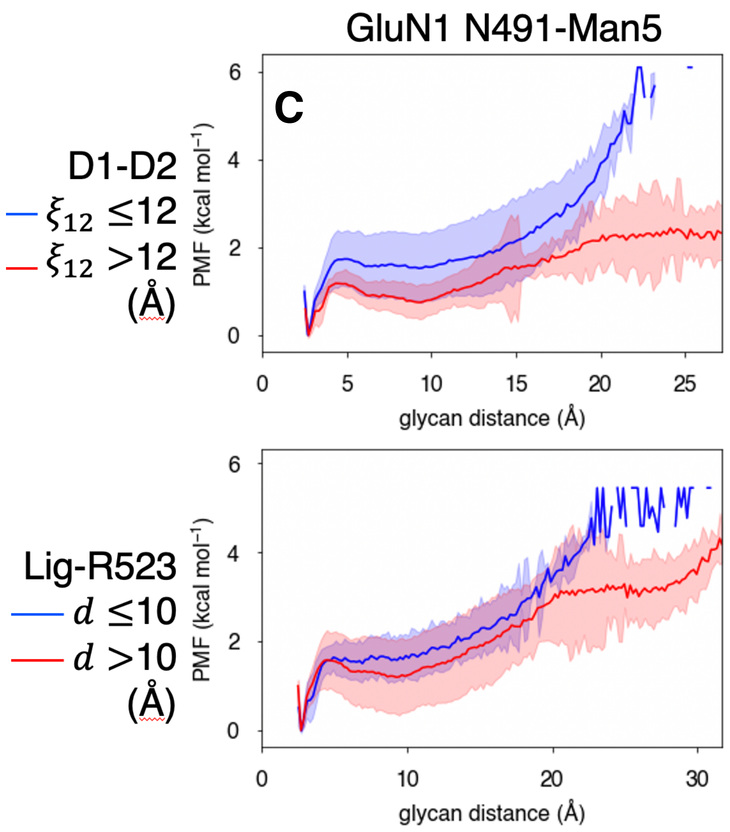


**Figure 6 – figure supplement 2.** Glycan-D2 distance dependence on LBD closure and agonist binding for the **(A)** GluN2A N443-Man5 glycan, **(B)** GluN2A N444-Man5 glycan, and **(C)** GluN1 N491-Man5 glycan.

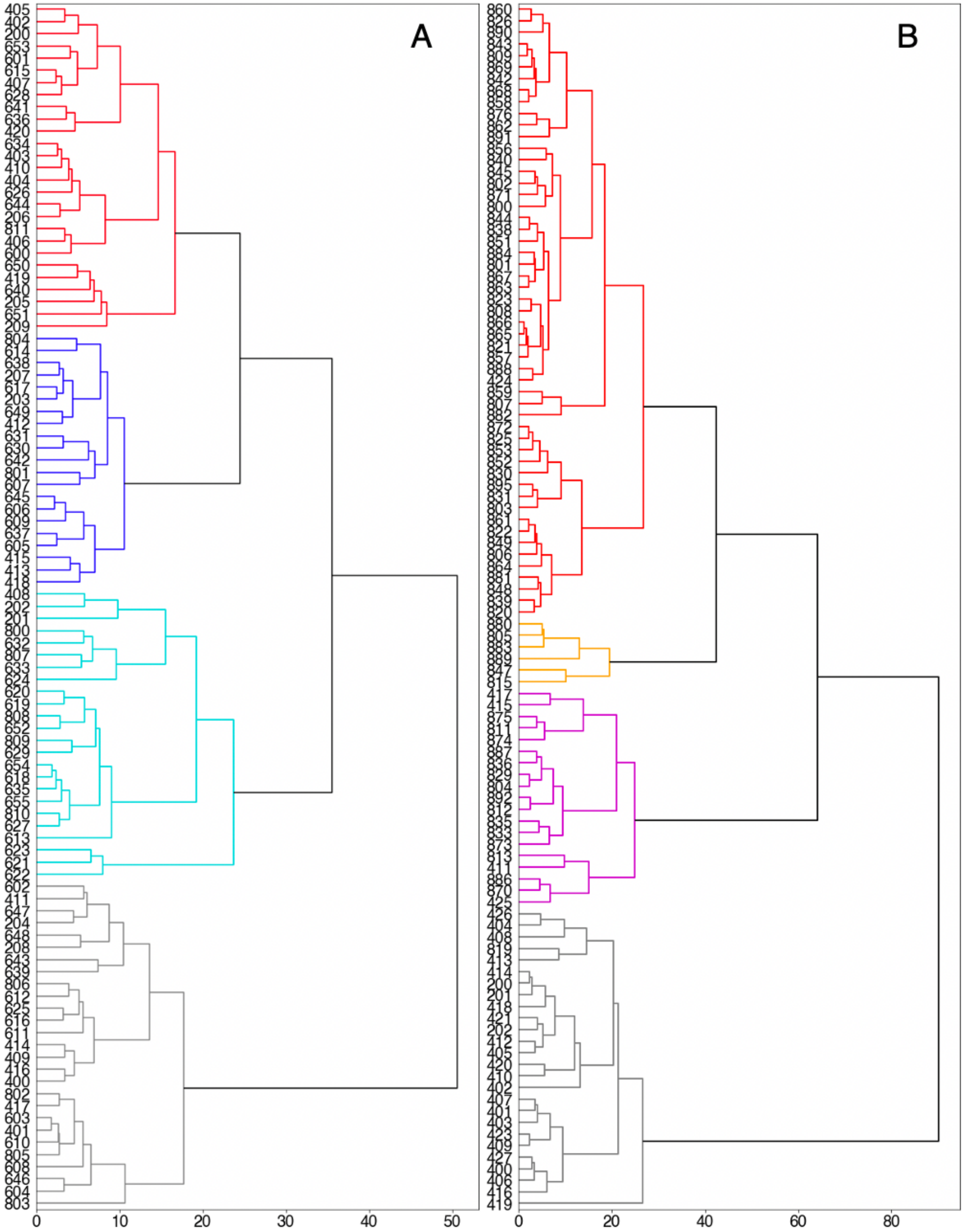


**Figures 1, 2 – figure supplement 4.** Dendrograms for hierarchical clustering of weighted average hausdorff distances for D-serine binding pathways for **(A)** GluN2A and **(B)** GluN1 according to the Ward linkage criterion.

**Legends for Datasets S1-S11. Data extracted from molecular dynamics simulations.**

**S1. 1_Simulation_Summary**: overview of simulation systems
**S2. 2_GluN2A**: record of all successful binding pathways in each simulation system for D-serine binding to GluN2A.
**S3. 3_Glu**: record of all successful binding pathways in each simulation system for glutamate binding to GluN2A.
**S4. 4_GluN1**: record of all successful binding pathways in each simulation system for D-serine binding to GluN1.
**S5. 5_contact_freq_N2_glyc**: per residue contact frequency analysis for D-serine binding to GluN2A by cluster identified with PSA.
**S6. 6_contact_freq_N1_glyc**: per residue contact frequency analysis for D-serine binding to GluN1 by cluster identified with PSA.
**S7. 7_Dser_vs_Glu**: comparison of relative residue contact frequency for D-serine and glutamate.
**S8. 8_bound_state_analysis**: per residue contact frequency analysis of the bound state for each agonist computed from lowest-energy conformers extracted from umbrella sampling simulations.
**S9. 9_conditional_prob_N2**: GluN2A residues most frequently contacted by D-serine given that the pathway results in successful binding – listed for each simulation system.
**S10. 10_conditional_prob_N1**: GluN1 residues most frequently contacted by D-serine given that the pathway results in successful binding – listed for each simulation system.
**S11. 11_glyc_vs_noglyc**: comparison of relative residue contact frequency during GluN2A and GluN1 binding pathways for glycosylated and non-glycosylated simulations.

**Legends for Movies S1-S2**

**Movie 1.** Process of D-serine (spheres) binding to the GluN2A LBD (light orange cartoon). D-serine diffuses in bulk solvent until it contacts LBD residues (sticks) that guide it into the binding site. Shown also are the GluN1 LBD (light blue cartoon) and N-linked Man5 glycans (sticks).
**Movie 2.** Process of D-serine (spheres) binding to the GluN1 LBD (light blue cartoon). D-serine diffuses in bulk solvent until it contacts LBD residues (sticks) that guide it into the binding site. Shown also are the GluN2A LBD (light orange cartoon) and N-linked Man5 glycans (sticks).
